# Genome-wide binding of posterior HOXA/D transcription factors reveals subgrouping and association with CTCF

**DOI:** 10.1101/073593

**Authors:** Ivana Jerković, Daniel M. Ibrahim, Guillaume Andrey, Stefan Haas, Peter Hansen, Catrin Janetzki, Irene González Navarrete, Peter N. Robinson, Jochen Hecht, Stefan Mundlos

## Abstract

Homeotic genes code for key transcription factors (HOX-TFs) that pattern the animal body plan. During embryonic development, Hox genes are expressed in overlapping patterns and function in a partially redundant manner. *In vitro* biochemical screens probing the HOX-TF sequence specificity revealed largely overlapping sequence preferences, indicating that co-factors might modulate the biological function of HOX-TFs. However, due to their overlapping expression pattern, high protein homology, and insufficiently specific antibodies, little is known about their genome-wide binding preferences. In order to overcome this problem, we virally expressed tagged versions of limb-expressed posterior Hox genes (*Hoxa9-13*, and *Hoxd9-13*) in primary mesenchymal limb progenitor cells (micromass). We determined the effect of each HOX-TF on cellular differentiation (chondrogenesis) and gene expression and found that groups of HOX-TFs induce distinct regulatory programs. We used ChIP-seq to determine their individual genome-wide binding profiles and identified between 12,540 and 27,466 binding sites for each of the nine HOX-TFs. Principal Component Analysis (PCA) of binding profiles revealed that the HOX-TFs are clustered in two subgroups (Group 1: HOXA/D9, HOXA/D10, HOXD12, and HOXA13 and Group 2: HOXA/D11 and HOXD13), which are characterized by differences in their sequence specificity and by the presence of cofactor motifs. Specifically, we identified CTCF binding sites in Group 1, indicating that this subgroup of HOX-proteins cooperates with CTcf. We confirmed this interaction by an independent biological assay (proximity ligation assay) and showed that CTCF is a novel HOX cofactor that specifically associates with Group 1 HOX-TFs, pointing towards a possible interplay between HOX-TFs and chromatin architecture.

## Introduction

The homeotic genes (*Hox* genes) are key regulators of development. They encode homeodomain transcription factors (HOX-TFs) that are expressed in an overlapping fashion along the anterior-posterior axis in all metazoans (McGinnis and Krumlauf 1992). In the vertebrate genome, *Hox* genes are organized in clusters with their order reflecting not only their expression along the anterior-posterior body axis but also their temporal expression (spatio-temporal collinearity). In most vertebrates, two rounds of whole-genome duplication have resulted in four clusters of *Hox* genes, coding for a total of HOX-TFs. All HOX-show high levels of sequence conservation between paralog groups (e.g. HOXA9 and HOXD9) and to a lesser extent between genes of the same cluster (e.g. HOXA1 to HOXA13) (reviewed in Gehring et al. 2009; Rezsohazy et al. 2015).

In the developing vertebrate limb, the posterior genes of the *HoxA* and *HoxD* clusters ( *HOX9-13*) are expressed along the proximo-distal axis following a collinear strategy (Zakany and Duboule 2007). Genetic experiments inactivating individual *Hox* genes revealed a remarkable redundancy within paralog groups controlling the development of the proximal (stylopod), middle (zeugopod), and distal (autopod) parts of the limb (Boulet and Capecchi 2004; Zakany et al. 2004). For example, neither the homozygous deletion of *Hoxa11* nor *Hoxd11* in mice leads to substantial malformations of the stylo-, or zeugopods. However, deletion of both *Hoxa11* and *Hoxd11* causes a severe truncation of the stylopod and loss of the zeugopod (Davis et al. 1995; Raines et al. 2015). A similar redundancy is observed between genes of the same cluster. Deletions, in mice, that encompass the entire *Hoxd13* gene cause the adjacent *Hoxd12* to be expressed in a *Hoxd13*-like pattern associated with the functional rescue of the *Hoxd13* deficiency. A similar deletion, removing *Hoxd13* and *Hoxd12* causes *Hoxd11* to be expressed in a *Hoxd13*-like pattern; however, *Hoxd11* is not able to rescue the loss of its two adjacent paralogs (Kmita et al. 2002).

In spite of the insights gained by these elegant series of genetic experiments, the high degree of *Hox* protein similarity and the overlap of expression domains have hindered the elucidation of the individual HOX-TF functions. HOX-TFs were also included in large biochemical surveys to identify the specific binding sequence of transcription factors (Berger et al. 2008; Jolma et al. 2013; Jolma et al. 2015). Two complementary studies applying 89 protein binding microarrays (PBM) and SELEX-seq on purified DNA-binding domains demonstrated that all posterior HOX-TFs bind to similar AT-rich sequences that vary in their 3”5’” region/” but share a characteristic TAAA sequence in their 3’ half. Moreover, two NMR based studies showed binding of HOXA13 to the HOXD13 site and *vice versa* (Zhang et al. 2011; Turner et al. 2015). Thus, the DNA binding specificity is not sufficient to explain individual HOX-TF function. More recent studies revealed a crucial role for cofactors in HOX-TF specificity. HOX-cofactors were shown to specifically alter the recognition sequence of the HOX-TFs by forming heterodimers (Joshi et al. 2007; Slattery et al. 2011; Jolma et al. 2015). Moreover, the analysis of HOX-cofactor specific binding sites suggested that these altered binding sites might be functionally more relevant for HOX binding than the HOX-TFs binding sites themselves (Crocker et al. 2015). However, due to high sequence homology, inadequate antibody specificity, and overlapping expression patterns little is known about genomic binding of the different HOX-TFs and how this might relate to their biological function.

Here, we have analysed and systematically compared the effects of nine limb bud-expressed HOX-TFs (HOXA9-13 and HOXD9-13) on cell differentiation and gene regulation and compare their genome-wide binding characteristics. To mimic the natural HOX environment as closely as possible, we used mesenchymal chicken limb bud cells and mild retroviral overexpression (Ibrahim et al. 2013). In this primary cell culture system (chicken micromass, chMM) the cells normally undergo chondrogenic differentiation; a process that can be altered by virally expressed transgenes (Ibrahim et al. 2013). Given the identical cell origin, culture conditions, and antibody use, this system allowed us to assess the distinctive properties of each HOX-TF and compare them to each other.

We find that certain HOXA/HOXD paralog TFs have opposing effects on chondrogenic differentiation and induce distinct regulatory programs in transduced cells. Further, by comparing the genome-wide DNA binding of nine HOX-TFs, in this experimental setting, we find that the posterior HOX-TFs can be separated into two groups (Group 1 and Group 2), with distinct binding motifs and distinct associations with cofactors. Finally, we characterized CTCF (the CCCTC-binding factor) as a novel cofactor of HOX-TFs and show that Group 1 but not Group 2 HOX-TFs bind thousands of CTCF-occupied sites in the chicken genome.

## Results

### Posterior HOX-TFs have distinct effects on gene regulation and differentiation of mesenchymal limb bud cells

To systematically compare the function of posterior HOX-TFs, we virally expressed FLAG-tagged versions of each TF in chicken micromass (chMM) cultures. First, we assessed the effect induced by the different HOX-TFs on chMM cultures. We noticed that some HOX-TFs promoted chondrogenic differentiation (HOXA9, HOXA10, HOXD10), while others inhibited the process (HOXD9, HOXD11, HOXA11, HOXD12, HOXA13, and HOXD13) (Figure 1A).

**Figure 1.**
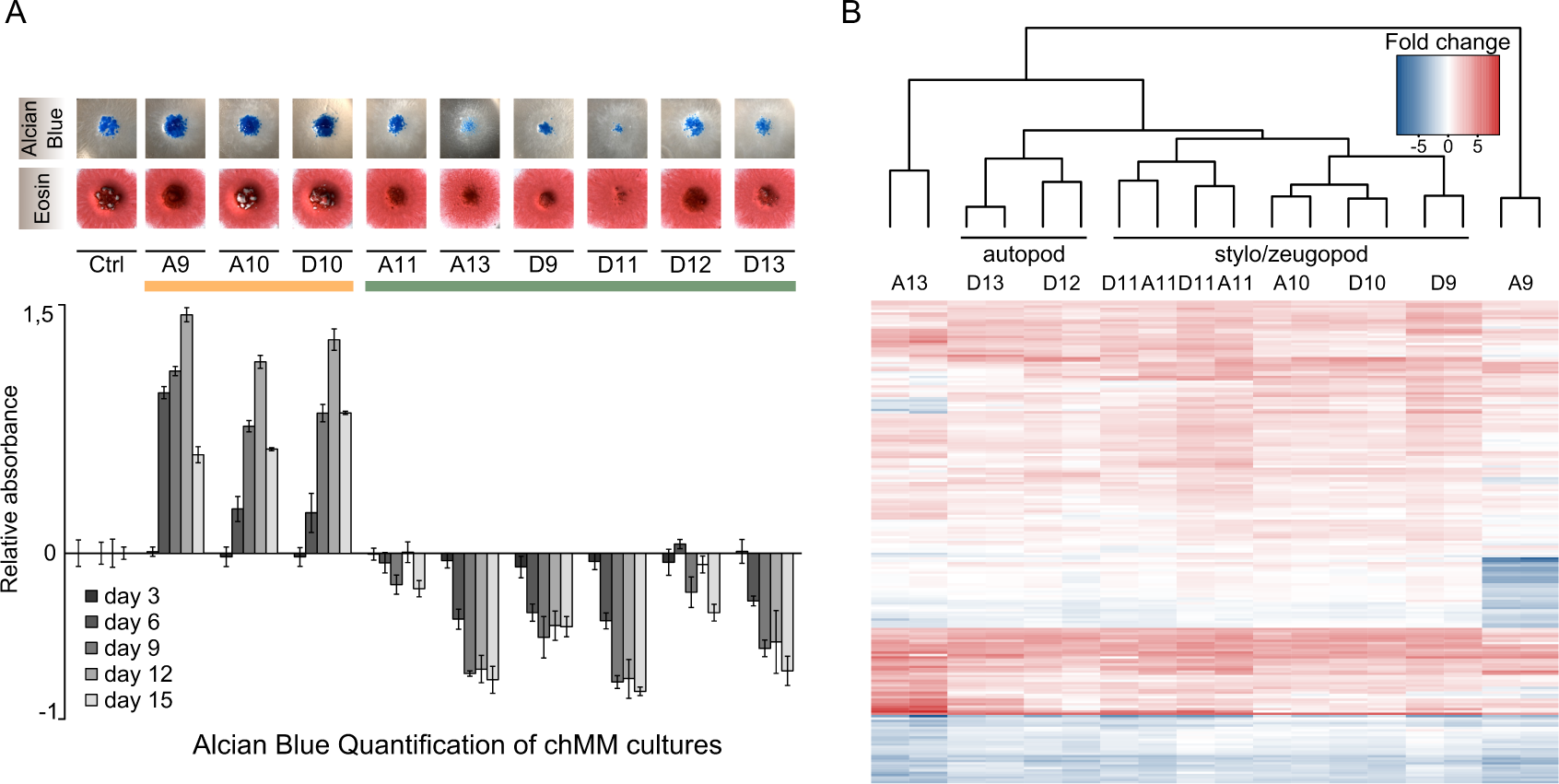
Viral expression of HOX-TFs in chicken micromass cultures (chMM) modifieschondrogenic cell differentiation. A) Individual HOX-TF expressing chMM cultures stained with Alcian blue (top) and Eosin (bottom)Alcian Blue staining of four biological replicates was quantified and compared to mock-infected chMM. Error bars indicate standard deviation from four replicates. B) Hierarchical clustering of differentially regulated genes in the nine HOX-TF expressing cultures (al RNA-seq shown in replicates). The top 50 differentially regulated genes from each sample were selected (Criteria: p-val ≤10e-5, base mean≥30, fold change≥2) and for each replicate the log2-transformed fold changes relative to mock-infected cultures of these genes were subjected to hierarchical clustering.

Interestingly, paralogue HOX-TFs did not always have the same general impact on the chondrogenic differentiation of the chMM. While *HOXA9* stimulated chondrogenic differentiation, its paralog *HOXD9* inhibited the same process. In contrast, *HOXA10* and*HOXD10* both promoted chondrogenic differentiation. *HOXA11* and *HOXD11* both inhibited chondrogenic differentiation, but to a very different extent. Finally, *HOXD13* and *HOXA13* both strongly inhibited cartilage formation; however, Eosin staining showed that the cell morphology of the *HOXA13*-expressing chMM was quite distinct from *HOXD12* or *HOXD13* cultures (Figure 1A).

The simple readout of the chMM morphology showed that the HOX-TFs induce distinct effects on cell differentiation. In order to comprehensively compare the effects on gene expression, we performed RNA-seq of HOX-TF expressing chMM cultures. We used DEseq2 (Love et al. 2014) to generate a list of genes that were differentially regulated compared with mock-infected chMM cultures. We then used the genes that were found among the 50 most strongly regulated genes in any of the nine datasets for hierarchical clustering (Figure 1B, Supplemental Table 1).

The hierarchical clustering recapitulated some of the main differences found between HOX-TFs that were detected in chMM gross morphology. HOX10 and HOX11 paralogs clustered together, while HOX9 paralogs, which bore striking differences in chMM morphology, clustered apart. Furthermore, the clustering process classified the paralog groups in an order that partially corresponded to their known role in limb development. The clustering separated the stylo-/zeugopod expressed HOX-TFs (*HOXD9, HOX10*/*11*) from the autopod expressed *HOXD12*/*13*. Two factors, *HOXA9* and *HOXA13*, clustered separately from all other HOX-TFs, indicating that the regulatory programs these factors induce are distinct from the other posterior HOX-TFs. Moreover, the *HOX11* paralogs induced transcriptional programs 154 so similar to one another that the clustering algorithm was not able to separate the two replicate datasets from each factor. Interestingly, two genes coding for subunits of the AP1 class of transcription factors, *JUN* and *FOS,* were among the most strongly upregulated genes in all of the datasets, suggesting that they might be direct targets of HOX-TFs. Ou analysis shows that, despite high homology and functional redundancy *in vivo*, the direct effects of paralog HOX-TFs in chMM cultures are distinct. While some can be similar (*HOXA10*/*HOXD10* and *HOXA11* /*HOXD11*) others can have opposing effects (*HOXA13*/*HOXD13* and *HOXA9*/*HOXD9*).

### Genome-wide binding reveals two distinct groups of HOX-TFs

We next wanted to assess whether analogous differences could be observed between paralog groups in their genome-wide binding preferences. We generated ChIP-seq profiles of the virally expressed HOX-TFs in chMM cultures using the αFLAG antibody. We identified between 12,540 and 27,466 binding sites for each of the nine HOX-TFs (Figure 2). We first assessed the binding sites shared between HOX-TFs from the same paralog groups by taking the 10,000 strongest peaks for each factor and then calculated the pairwise overlap between all HOX-TFs. Similar to the results of the expression analysis, the HOX10 and HOX11 paralogs shared more peaks (78-81% and 85-86%, respectively) than the HOX9 and HOX13 paralogs (65-60% and 24-16%) (Supplemental Figure 1A).

**Figure 2.**
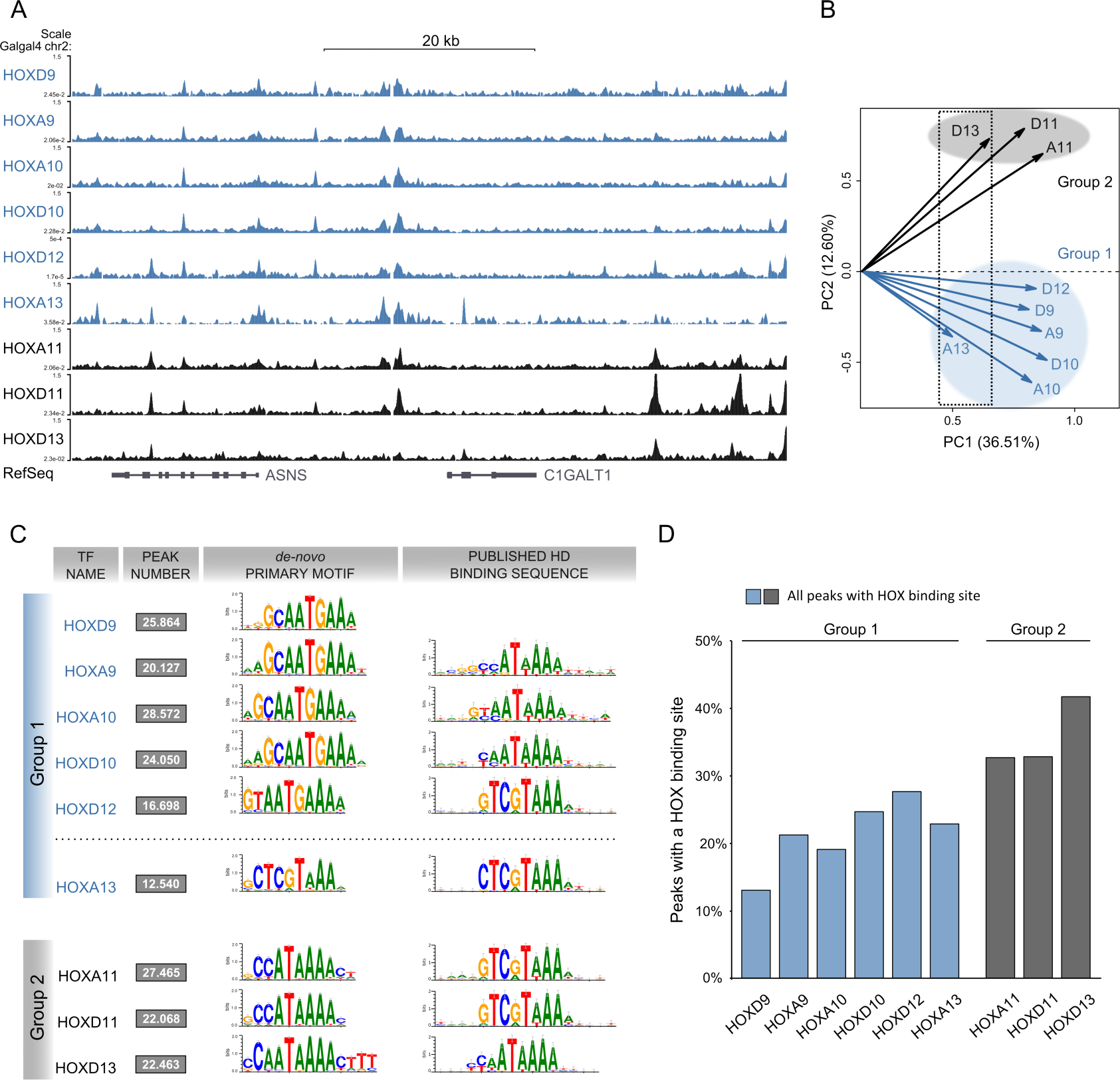
Genome-wide binding profiles of posterior HOX-TFs reveals two groups of binding. A) ChIP-seq profiles of nine posterior HOX-TFs (Group 1 – blue, Group 2 – black). B) Principal Component Analysis (PCA) analysis of HOX-TF peaks. HOX13 paralogs cluster separate on PC1 (dotted rectangle). PC2 reveals two distinct groups of HOX-TFS, Group 1 (blue) and Group 2(black). C) *De novo* motif analysis for the HOX-TFs. Primary motifs obtained from the top 5,000 peaks in comparison to the previously identified motifs for their respective homeodomains (Berger et al., 2008). Group 1 sequence preferences (except HOXA13) are distinct from Group 2. See Supplemental Fig. 1B for additional HOX-like motifs identified in the Top 5,000 peaks. D) Quantification of peaks carrying binding sites (sequences matching any of the top 3 HOX-TF motifs; FIMO p value≤ 0.0001). Each peak carrying a sequence match is counted only once. Binding site count in top 1,000 and top 10,000 peaks are shown in Supplemental Figure 1C.

Next, we performed a principal components analysis (PCA) to compare the datasets in an unbiased way, using the identified peaks as input (Figure 2B). PCA showed that the binding of HOX-TF paralogs seemed to be more similar than their effects on chMM differentiation and gene expression. HOXA13 and HOXD13 were a notable exception as they clustered separately from the other HOX-TFs along PC1 (Figure 2B, dashed box). In addition, they were also very different from one another in PC2. A comparison of all tested HOX-TFs in PC2 revealed a surprising separation into two groups, which neither reflected the effects on cell differentiation and gene expression, nor the sequence homology of the TFs. Group 1 comprised HOXA/D9, HOXA/D10, HOXD12, and HOXA13 (Figure 2B, blue) and Group 2 comprised HOXA/D11 and HOXD13 (Figure 2B, black).

To find a possible cause for this separation, we first tested whether the grouping could be attributed to the sequence-specificity of the TFs. For this we performed *de novo* motif analysis using the peak-motifs algorithm (Medina-Rivera et al. 2015) with the 5,000 strongest peaks as input and compared it to the published results from PBM and SELEX-seq (Figure 2C and Supplemental Figure 1B). This comparison showed a general similarity between *in vitro* and ChIP-seq derived motifs. However, several sequence features had not been detected in the previously published datasets. We found a prominent G at the 5’ end of all Group 1 motifs (HOXA/D9, HOXA/D10, HOXD12, and HOXA13), which had also been detected using SELEX-seq (Jolma et al. 2013). More striking, we found that the TAAA 3’ end, which is a characteristic of posterior HOX-TFs, changed to a TGAAA in all Group 1 HOX-TFs, with the notable exception of HOXA13.

The motifs identified for HOXA13 and HOXD13 were identical to the ones detected in PBM/SELEX-seq. In contrast, the primary motif of HOXA11 and HOXD11 did not overlap with those detected in the corresponding *in vitro* datasets. Specifically, the CCATAAA motif (HOXA/D11) we observed was highly similar to a change in sequence specificity that HOXA10 undergoes when co-binding with PBX4 (Jolma et al. 2015). Generally, motif analysis for the HOX-TFs identified not only primary motifs, but also several alternative HOX-like motifs, suggesting that the DNA-dependent binding of HOX-TFs might be less sequence-driven than other TFs (Supplemental Figure 1B).

Group 1 (HOXA/D9, HOXA/D10, HOXD12, and HOXA13) and Group 2 (HOXA/D11 and HOXD13) HOX-TFs also revealed differences, when we considered the fraction of ChIP-seq peaks that contained a HOX-TFs binding site (Figure 2D and Supplemental Figure 1C). The number of peaks carrying a HOX-binding site (i.e. matching one of the top three HOX motifs) was relatively low in general, ranging from as little as 15% (HOXD9) to 43% (HOXD13).Interestingly, the three Group 2 HOX-TFs had the highest number of HOX binding sites in contrast to the Group 1 HOX-TFs, which displayed the lowest number of peaks carrying HOX binding sites. To exclude the effect of weak and maybe indirect binding sites from the analysis, we performed the same analysis for the 10,000 and 1,000 strongest peaks(Supplemental Figure 1C). Although the fraction of binding site-containing peaks slightly increased, the general distribution stayed the same.

### *De novo* motif analysis finds putative HOX-cofactors

The relatively low numbers of HOX-TF peaks containing HOX binding sites indicated that other factors might contribute to DNA binding. Sequence analysis of ChIP-seq peaks allows not only for the detection of sequence-specific binding sites, but also for the identification of putative cofactors. Therefore, we performed a *de novo* motif analysis using all peaks as input and then compared the non-HOX like motifs to the literature and to large TF motif databases (JASPAR (Mathelier et al. 2015), footprint DB (Sebastian and Contreras-Moreira 2014))(Figure 3).

**Figure 3.**
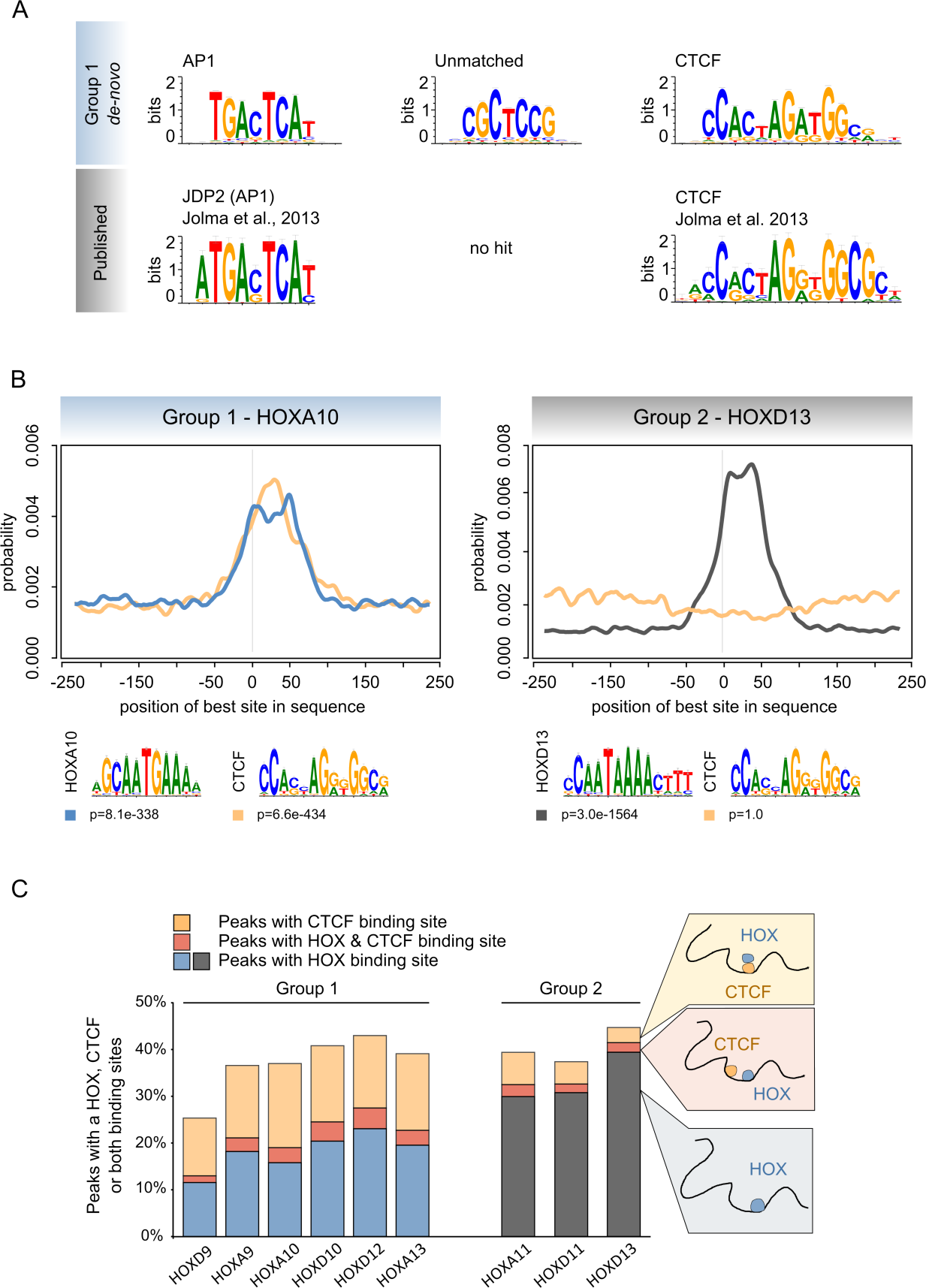
AP1 and CTCF binding sites are overrepresented in Group 1 HOX-TF binding sites. A) *De novo* motif analysis of all Group 1 HOX-TF peaks (here, HOXA10 results) identifies overrepresented binding sites. A comparison of these motifs to known AP1 and CTCF motifs is shown below. B) Centrimo analysis identifies the position of best binding site matches in all peak sequences. Blue and black lines indicate enrichment of the given HOXA10 or HOXD13 motif shown below, respectively.Yellow lines indicate enrichment for CTCF motif shown below. CTCF binding sites are centrally enriched in HOXA10 peaks. C) The overlap of peaks containing a HOX (Group 1-blue, Group 2-black) or a CTCF (yellow) binding site. The red overlap indicates peaks containing a HOX and a CTCF binding site.

In Group 2 of HOX-TFs, we were not able to detect any clear cofactor motif. In contrast, we found three putative cofactor motifs in five out of the six Group 1 HOX-TF peak sets. The first motif was the well-characterized TGANTCA AP1 binding site (Glover and Harrison 1995)(Figure 3A). A second motif, CGCTCCG was detected with high specificity in the HOXA9 and HOXD9 peaks and with lower specificity (but still among the top 5) in the HOXA10, HOXD10 and HOXD12 peaks (Supplemental Figure 2A and 2B). This motif was particularly enriched in HOXA13 peaks (Supplemental Figure 5). We were not able to find matching or similar motifs in the JASPAR and footprint-DB databases, raising the possibility that it either represented the binding site of an uncharacterized TF or a composite binding site recognized by a dimerized TF complex. As a third motif, we detected a 12bp long GC-rich motif in all Group 1HOX-TF datasets except HOXA13. This motif perfectly matched the known motif of the CCCTC-binding factor (CTCF), a well described TF involved in gene regulation and genome architecture (Figure 3A and Supplemental Figure 2)(Barski et al. 2007).

The *de novo* discovery of cofactor motifs can be masked by the strong overrepresentation of the primary motif. To exclude this possibility, we performed a reverse search and identified and counted all matches to CTCF (Figure 3, Supplemental Figure 3) or AP1 (Supplemental Figure 4) binding sites in the nine Hox-TF data sets. For the CTCF binding sites, this reverse search revealed a characteristic difference between Group 1 and Group 2 HOX-TFs. Altogether, 12-21% of all Group 1 HOX-TF peaks, but only 4-9% of Group 2 HOX-TF peaks contained a CTCF binding site (Supplemental Figure 3A). In contrast, we identified AP1 binding sites in about 3-6% of all peaks of the different HOX-TFs and there seemed to be no distinction between Group 1 and Group 2 HOX-TFs (Supplemental Figure 4).

Next/”we mapped the position of the CTCF binding sites within the HOX-TF peaks and found that in Group 1, but not Group 2, the CTCF sites were located predominantly near the peak summits (Figure 3B and Supplemental Figure 3B), suggesting a binding mode in which the HOX-TF binds indirectly via CTcf. This was further supported by a discriminatory motif analysis, which revealed that Group 1 HOX-TF peaks contained either a HOX or a CTCF binding site and that only a minority of HOX-TF peaks contained binding sites for both TFs (Figure 3C).

### Group 1 HOX-TFs and CTCF/cohesin co-bind genome-wide

Motif analysis indicated that CTCF and Group 1 HOX-TFs might co-bind to many sites throughout the genome. We therefore mapped CTCF binding sites genome-wide by virally expressing FLAG-tagged CTCF in chMM cultures (Figure 4) and performed ChIP-seq using the αFLAG antibody. From the same sample, we also performed ChIP-seq for endogenous RAD21, a subunit of the cohesin complex and an important CTCF-cofactor (Faure et al. 2012).We identified 22,357 CTCF and 17,589 RAD21 binding sites. Similar to previous reports, CTCF and RAD21 co-bound to 53% of all CTCF and to 67% of all RAD21 peaks. We then tested how many HOX-TF peaks overlapped with CTCF or RAD21 peaks. We observed that the characteristic distinction between Group 1 and Group 2 HOX-TFs could be recapitulated at ChIP-seq binding sites. Indeed, Group 1 HOX-TFs shared between 15% and 24% of their peaks with CTCF (12-20% with RAD21), whereas only 3-8% of Group 2 peaks overlapped with CTCF (3-7% with RAD21) (Figure 4C and Supplemental Figure 6A, B and C).

**Figure 4.**
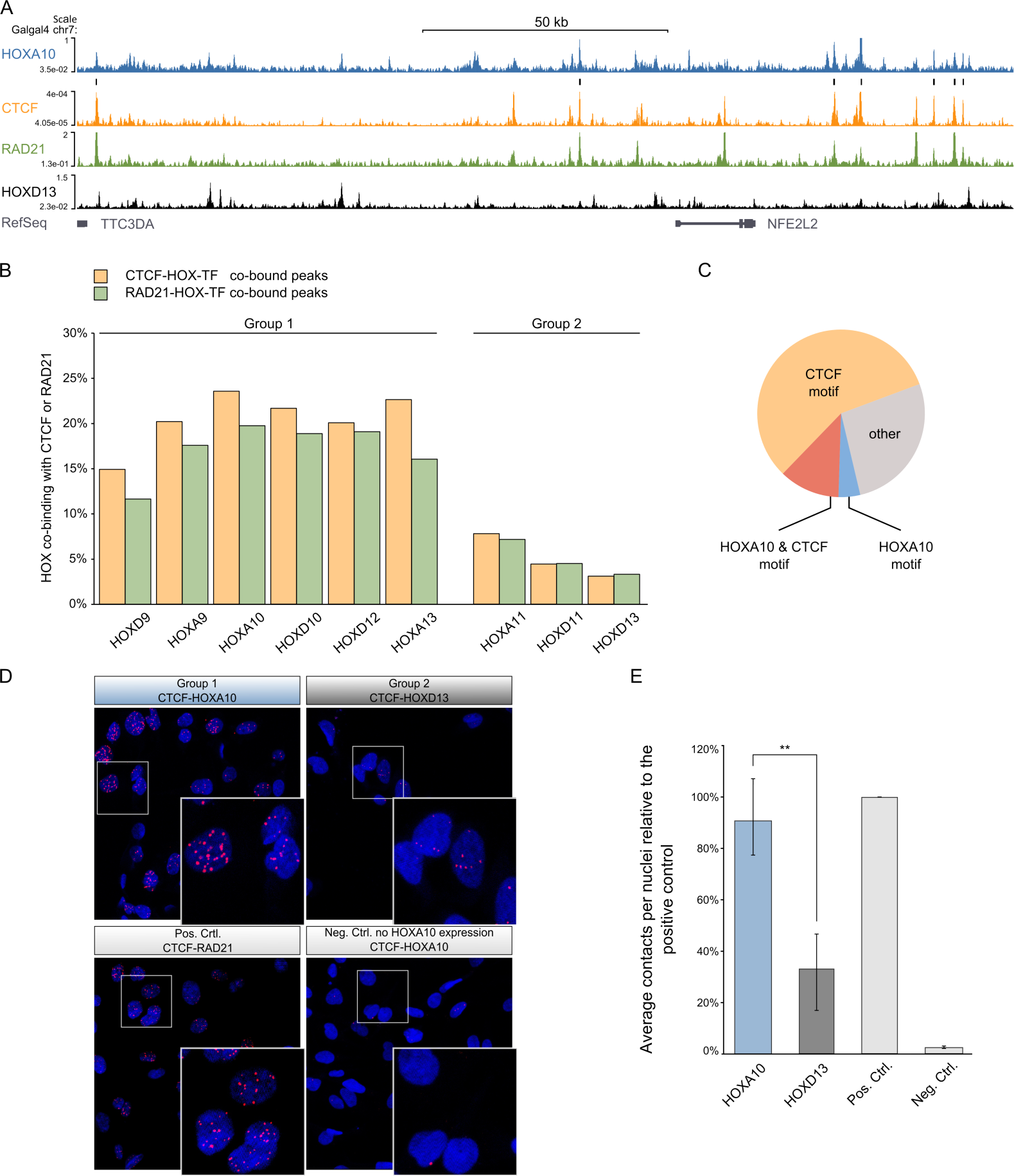
Group 1 HOX-TFs and CTCF/RAD21 share thousands of binding sites throughout the genome. A) ChIP-seq tracks of HOXA10, CTCF, RAD21, and HOXD13. Black bars above the CTCF-track indicate HOXA10/CTCF co-bound sites. B) Percentage of HOX-TF peaks overlapping with CTCF (yellow) or RAD21 (green) peaks. C) Presence of HOXA10 and CTCF binding sites in the HOXA10-CTCF co-bound peaks. D, E) Proximity Ligation Assay in DF1 chicken fibroblasts. D) Top row: DF1 cells expressing 3xFLAG-HOXA10 (left) or 3 x FLAG-HOXD13 (right). PLA was performed using αFLAG and αCTCF antibodies.Bottom row: Positive Control (left) shows HA-CTCF expressing DF1 cells. PLA was performed with αHA and αRAD21. Negative Control (right) shows non-transfected DF1 cells, PLA performed with αFLAG and αCTcf. E) Quantification of PLA experiments. Contacts were counted with ImageJ and divided by the number of nuclei in three independent biological replicates (see Supplemental Figure 7). The graph shows the percentage of counted contacts relative to the positive control. The standard error of the mean is shown for every sample. A T-test was performed to measure the significance of the contact difference between HOXA10 and HOXD13 (Student’s T p-value<0.005).

Finally, we investigated the presence of CTCF and HOX binding motifs at shared binding sites with HOXA10 (Group 1) as a representative example. We looked for underlying binding sites in the 24% of HOXA10 peaks that are shared with CTCF and observed that 69% of them contained a CTCF binding site (23% in all HOXA10 peaks). In contrast, only 16% of the peaks had a HOXA10 binding site (18% in all HOXA10 peaks), suggesting that HOXA10 indirectly binds to these CTCF-shared peaks via CTcf. Taken together, motif analysis of HOX-TF binding sites and ChIP-seq for CTCF/RAD21 both found Group 1, but not Group 2 HOX-TF binding associated with CTCF/cohesion.

### Group 1 HOX-TFs and CTCF interact in the nucleus

Both, motif analysis and peak overlap strongly suggested an interaction between Group 1 HOX-TFs and CTcf. To test this possibility, we made use of the proximity ligation assay (PLA)(Soderberg et al. 2006). The PLA assay allowed us to assess protein-protein interactions *in situ,* in a quantifiable and sensitive manner. We expressed FLAG-tagged *HOXA10* (Group 1) in chicken DF1 cells and performed the PLA assay using αFLAG antibody and an endogenous αCTCF antibody. We readily detected CTCF-HOXA10 interaction in the nucleus that was almost as strong as the interaction of CTCF with RAD21, which we used as a positive control (Figure 4). We also performed the same assay with CTCF and the Group 2 HOXD13 protein, for which our ChIP-seq data had predicted a weaker interaction. In this case we measured a signal above our negative control (DF1 cells expressing CTCF alone), but less than for the CTCF-HOXA10 interaction (Figure 4D and E and Supplemental Figure 7).

## Discussion

In this study, we systematically compared the effect of nine limb-bud expressed HOX-TFs on the differentiation and gene regulation of primary mesenchymal limb bud cells. Hierarchical clustering of the regulated genes delineated two groups of HOX-TFs: HOX9/10/11,and HOXD12/HOXD13 that, during limb development, are expressed in the stylo/zeugopod and autopod, respectively. The distinction between these two groups is in accordance with genetic experiments in mice demonstrating that *Hoxd12*, but not *Hoxd11* is able to substitute for a loss of *Hoxd13*(Kmita et al. 2002). Another interesting observation was that *HOXA9* and *HOXA13* clustered separately from the other factors. Differences between *HOX9* and *HOX13* paralogs, in contrast to the more similar *HOX10* and *HOX11* paralogs, were also apparent in their distinct effects on chMM differentiation. This divergence might be explained by the fact that the *HOX9* and *HOX13* paralog groups are the only posterior HOX-TFs which retained all four copies of the genes, thereby reducing the selective pressure on each paralog and allowing their neo-functionalization (Gehring et al. 2009).

Our systematic comparison focused on the effects of individual HOX-TFs and their genome-wide binding. However, HOX-TFs are rarely expressed alone *in vivo*, but are rather co-expressed in overlapping patterns and exert their specific function in this biochemical 306 context. Although HOX-TFs induced distinct effects in our experiments, their combinatorial or antagonistic action *in vivo* might play an important role in the developing embryo.Investigation of the *in vitro* sequence specificity of individual HOX-TFs showed that their homeodomains bind largely similar sequences (Berger et al. 2008; Jolma et al. 2013). Subsequent studies, however, revealed that the binding of cofactors changes the original HOX binding site resulting in recognition sites that are markedly different (Slattery et al. 2011; Crocker et al. 2015; Jolma et al. 2015). Both observations are reflected in the results of our ChIP-seq experiments. The low number of direct binding sites in HOX-TF peaks found in our experiments is in concordance with results from *Drosophila*, where low-affinity binding sites for the HOX-TF Ultrabithorax (Ubx) in complex with its cofactor Extradenticle (Exd) were shown to be biologically more significant (Crocker et al. 2015). Our analysis also highlights 317 the role cofactors play in directing HOX-TF binding. The primary motif for both HOX11 paralogs was in many ways different from the *in vitro* determined monomer specificity and rather revealed a composite binding site like the one bound by a HOXA10-PBX4 dimer (Jolma et al. 2015). Furthermore, our data indicate a relationship between HOX-TFs and the AP1 class of TFs. AP1 binding sites were found in 5% of all HOX-TF peaks and *JUN* and *FOS* were also strongly upregulated by all HOX-TFs, suggesting a mechanism of cofactor cross-regulation. To our knowledge, AP1 has not been linked to limb patterning or HOX-TFs.However, these factors are known to be involved in a wide array of developmental and cell differentiation processes (Hess et al. 2004) and our results suggest AP1 may potentially have a role in mediating *HOX*-driven limb patterning.

PCA analysis separated the HOX-TF binding sites in two subgroups along PC2. We tried to identify the underlying cause for this distinction between HOX-TF binding sites and found co-binding with CTCF to correlate with Group 1 HOX-TF binding. We also describe CTCF as a novel cofactor of Group 1 HOX-TFs. CTCF/cohesin are now well-established factors with important functions in the spatial organization of the genome into topologically associating domains (TADs) (Dixon et al. 2012; Zuin et al. 2014; Sanborn et al. 2015). Among other functions, they have been shown to directly mediate enhancer-promoter contacts (Faure et al. 2012; Merkenschlager and Odom 2013; Ing-Simmons et al. 2015). The co-occupancy of CTCF/cohesin and HOX-TFs throughout the genome points to a possible role for this type of developmental TF in enhancer-promoter communication and beyond. In fact, HOX13 TFs have recently been implicated as regulators at the *HoxD* locus, where two adjacent TADs regulate the gene expression in the proximal and distal limb, respectively (Andrey et al. 2013; Beccari et al. 2016). Specifically, HOX13 proteins did not regulate individual enhancers, but rather restructured the chromatin architecture of the locus in a way so that contacts with one (the telomeric) TAD were repressed, whereas contacts with the other (centromeric)TAD were promoted (Beccari et al. 2016). A related observation was recently reported in *Drosphila* for CTCF/Cohesin and Smad-TFs, which are the transcriptional effectors of TGFß/BMP signalling (Van Bortle et al. 2015). The Smad-TFs co-localized in a CTCF-dependent manner to CTCF binding sites within TADs and might be involved in sculpting the TAD to enable transcriptional regulation. The observed connection of certain developmental TFs with CTCF/cohesin architectural proteins suggests an important fundamental regulatory role for HOX and other TFs that extends beyond the control of individual gene expression.

## Material and Methods

### Construction of viral vectors and chicken micromass cultures

*HOX* and *CTCF* coding sequences were amplified from chicken embryonic limb buds cDNA (HH27) and cloned into RCASBP-viruses as previously described (Ibrahim et al. 2013). DF1 cells were transfected in a 6 cm dish with 3 µg of each RCASBP(A) plasmid using Polyethylenimine (Polyscience Inc. #24765-2) and NaCl. Cells were expanded and stressed on starvation media whereupon the supernatant was harvested on three consecutive days. The supernatant was then centrifuged to produce the concentrated viral particles of high titer, 10^8^ viral particles/ml or higher. The infection of chMM cultures and the histological assessment were performed as described elsewhere (Ibrahim et al. 2013). For the quantification of chondrogenesis, cultures after 3, 6, 9, 12 and 15 days post infection were fixed and stained with Alcian Blue. After 2 washes with 1 x PBS, the quantification of incorporated Alcian Blue was determined by extraction with 6 M guanidine hydrochloride, followed by photometric measurement at A 595 nm.

### Chromatin preparation and ChIP-sequencing

Chromatin Immunoprecipitation was performed as described previously (Ibrahim et al. 2013), *all experiments were performed in two biological replicates (except HOXA13)*. Briefly, chMM cultures were harvested after 6 days of culture by adding digestion solution (0.1% collagenase (Sigma, #C9891) and 0.1% Trypsin in 1x PBS) to obtain a roughly single-cell suspension. Cells were taken up in 10 ml cold chMM (DMEM: HAMF11 with 10% FBS, 10% CS, 1% L-glutamine and 1% Penicilin-Streptomycin) medium and fixed for 10 min on ice with 1% formaldehyde. The extraction of nuclear lysate was performed as described in Lee et al. (2006) and chromatin was sonicated with a Diagenode Bioruptor (45 cycles - 30-sec pulse, 30-sec pause, HI power). For ChIP, 25-35 µg of chromatin was incubated with 6-8 µg of 544 antibody overnight. The next day blocked magnetic beads were added and incubated 545 overnight, followed by 6 washes with RIPA and one with TE (Lee et al. 2006). After elution, 546 the preparation of the library for pulled down DNA was performed as described previously (Ibrahim et al. 2013).

### RNA&sequencing

Cells from harvested chMM cultures were separated prior to fixation of the ChIP samples and RNA was isolated from these cells using an RNaeasy Qiagen kit. RNA-seq libraries were 552 constructed as described previously (Ibrahim et al. 2013), by selecting for fragment sizes between 300-bp and sequenced single-end 50 bp using Illumina technology.

### Proximity Ligation Assay (PLA)

DF1 cells were transfected with RCASBP(A)-3x FLAG-HOXA10, RCASBP(A)-3x FLAG-HOXD13, or RCASBP(C)-HA-CTCF, respectively. The cells were cultured for at least 6 days to ensure a high cellular infection rate. Upon confluency cells were transferred to 10 mm cover slips and 559 further incubated for one day. Cells were fixed for 10 min with 4% PFA, blocked with TSA (10% horse serum, 0,5% PerkinElmer blocking reagent [#FP1020] and 0.01% Triton-X-100 in 1x DPBS) and incubated with appropriate primary antibodies (in 10% horse serum in 1x DPBST) overnight at 4°C. Primary antibody combinations were: 1) FLAG-HOX and CTCF 563 interaction: m-αFLAG M2 and rb-αCTCF; and 2) HA-CTCF and RAD21 interaction: m-αHA and rb-αRAD21. Antibody concentration for PLA were tested and used as follows: m-αFLAG M2 1:20000 (Sigma, F1804), m-αHA.11 1:8000 (BioLegends, #901501), rb-αCTCF 1:20000 (ActiveMotif, #61311) and rb-αRAD21 1:1000 (Abcam, ab992).

After primary antibody incubation, the PLA assay was performed using the Duolink In Situ Fluorescence Kit (Sigma, #DUO92101-1KT) according to manufacturer’s instructions. Protein-protein interactions were analyzed by using confocal imaging on a Zeiss LSM700 and the Axiovert Zen software.

For the quantification of PLA experiments, the contacts in several independent frames were counted using ImageJ and divided by the number of nuclei in the frame. The PLA experiments were performed in at least two independent experiments.

## Bioinformatic Analyses

### ChIP-seq

#### Processing and peak analysis

Quality filtering and read mapping were performed as described previously (Ibrahim et al. 2013). Reads were mapped against the g*alGal4* reference genome. Reproducible peaks were identified using the MASC2 (Zhang et al. 2008) peak caller and IDR pipeline (Kundaje 2012; Landt et al. 2012). For calculating peak overlaps we used bedtools (Quinlan and Hall 2010). Summits were extended +/-150 bp and two peaks were considered overlapping if the overlap was >= 100 bp. The Principal Components Analysis was performed on detected reproducible peaks as described previously (Ibrahim et al. 2013).

#### Sequence analysis and peak overlap

*De novo* motif analysis was performed using the peak-motifs algorithm (Medina-Rivera et al. 2015) and the sequences +/-75 bp surrounding the respective peak summits. For counting individual binding sites in the peaks we extracted the sequences +/-150bp surrounding the peak summit. Next, the position weight matrix (PWM) of the top three motifs that described the HOX binding site were used together with the FIMO software (Grant et al. 2011) to obtain the peaks containing a binding site (p < 0.0001). Following this, all peaks that had a sequence match for any of the three motifs were counted as carrying a binding site. A trimmed version of the CTCF-matrix according to the (Barski et al. 2007)(see Fig. 2) was used for counting the occurrences of CTCF binding sites,. For Centrimo (Bailey and Machanick 2012) analysis, we used sequences +/-250 bp of the peak summit and trimmed versions of the PWMs as seen in the motif logos.

### RNA-seq analysis

RNA-sequencing reads were mapped to the chicken reference genome galGal4 using the STAR mapper (Dobin et al. 2012) (splice junctions were based on RefSeq/ENSEMBL gene annotations; options included: alignIntronMin 20, alignIntronMax 500000, outFilterMultimapNmax 5, outFilterMismatchNmax 10, and --outFilterMismatchNoverLmax 0.1). Read counts for individual genes were generated for a gene list combining the RefSeq (galGal4) and ENSEMBL (release 75) gene annotations.

Log2 Fold changes for differential expression were calculated using DEseq2 (Love et al. 2014). The top 50 regulated genes were filtered according to p-value < 10^-^5, a minimum base mean >30 and a fold change >2. For hierarchical clustering, all genes were included that were among the top 50 regulated genes in at least one of the datasets. The log2-transformed fold changes, as compared with control cultures, were then used as the input for the R heatmap3 hierarchical clustering algorithm.

## Acknowledgements

IJ was supported by a PhD stipend from the Berlin-Brandenburg School of Regenerative Therapies. This study was supported by a grant from the Bundesministerium für Bildung und Forschung (Förderkennziffer FKZ 1315848A) to SM, JH, and PNR. We thank Christina Paliou, Franziska Martinez Real, Martin Franke and other members of the Mundlos laboratory for 618 their critical reading of the manuscript.

